# α-Melanocyte-Stimulating Hormone Regulates Pathological Cardiac Remodeling by Activating Melanocortin 5 Receptor in Cardiomyocytes

**DOI:** 10.1101/2023.04.06.535956

**Authors:** Anni Suominen, Guillem Saldo Rubio, Saku Ruohonen, Zoltán Szabó, Lotta Pohjolainen, Bishwa Ghimire, Suvi T. Ruohonen, Karla Saukkonen, Jani Ijas, Sini Skarp, Leena Kaikkonen, Minying Cai, Sharon L. Wardlaw, Heikki Ruskoaho, Virpi Talman, Eriika Savontaus, Risto Kerkelä, Petteri Rinne

## Abstract

**Background:** α-Melanocyte-stimulating hormone (α-MSH) regulates diverse physiological functions by activating melanocortin receptors (MC-R). α-MSH is predominantly expressed in the pituitary gland, but it is also found in several peripheral tissues such as the skin and heart. However, the role of α-MSH and its possible target receptors in the heart remain completely unknown. Therefore, we sought to investigate whether α-MSH could be involved in the regulation of pathological cardiac remodeling.

**Methods:** Tissue α-MSH concentrations and the effects of chronic α-MSH administration were investigated in mice subjected to transverse aortic constriction (TAC). Rat H9c2 cells, neonatal mouse ventricular myocytes and human induced pluripotent stem cell-derived cardiomyocytes (hiPSC-CM) were used to study the effects of α-MSH and selective MC-R agonists. Inducible cardiomyocyte-specific melanocortin 5 receptor (MC5-R) knockout mouse model was engineered to investigate the role of MC5-R in cardiac hypertrophy.

**Results:** α-MSH was highly expressed in the mouse heart, particularly in the ventricles, and its level was reduced in the left ventricles of TAC-operated mice. Administration of a stable α-MSH analogue protected mice against TAC-induced cardiac hypertrophy and systolic dysfunction. *In vitro* experiments revealed that cardiomyocytes serve as effector cells for the α-MSH mediated antihypertrophic signaling and that selective activation of MC5-R mimics the actions of α-MSH. In keeping with these findings, MC5-R was downregulated in the failing mouse heart and stressed hiPSC-CMs. Silencing of MC5-R in mouse cardiomyocytes induced hypertrophy and fibrosis markers *in vitro* and aggravated TAC-induced cardiac hypertrophy and fibrosis *in vivo*. Conversely, pharmacological activation of MC5-R improved systolic function and reduced cardiac fibrosis in TAC-operated mice.

**Conclusions:** α-MSH is expressed in the heart and protects against pathological cardiac remodeling by activating MC5-R in cardiomyocytes. These results suggest that analogues of naturally occurring α-MSH, that have been recently approved for clinical use and have agonistic activity at MC5-R, may be of benefit in treating heart failure.

## Introduction

Heart failure is one of the leading causes of hospitalization,^1-3^ causing enormous burden on our economic and health care systems. Despite currently available therapeutics and advancements in the clinical management of heart failure and its associated complications, many challenges still remain as morbidity and mortality rates are showing no signs of decrease,^2^ highlighting unmet medical need to develop novel disease-modifying therapies for this disease.

Heart failure is a common end-stage manifestation for many cardiovascular diseases and it primarily results from hypertension, ischemic heart disease and aortic valve stenosis. Hypertension, for instance, increases cardiac workload and induces left ventricular (LV) hypertrophy, which is compensatory and adaptive in nature and helps to maintain circulatory homeostasis. However, in the long term, the stressed heart undergoes maladaptive remodeling resulting in the development of fibrosis, LV dilatation and dysfunction, thus predisposing individuals to heart failure.^4,5^ Melanocortins are peptide hormones that are proteolytically cleaved from their precursor molecule known as proopiomelanocortin (POMC) and post-translationally modified into melanocyte stimulating hormones (α-, β- and γ-MSH) and adrenocorticotrophic hormone (ACTH).^6,7^ These peptide hormones mediate their biological actions through five different but closely related G-protein coupled melanocortin receptors (MC1-R to MC5-R) that are distributed in many tissues and involved in the regulation of important physiological functions including skin pigmentation, steroidogenesis, energy homeostasis, sexual function and exocrine secretion.^8^ Melanocortins have different affinities for their target receptors but α-MSH is unique in this regard as it has the ability to activate all MC-R subtypes except MC2-R (also known as ACTH receptor).^8,9^ Classical physiological effects of α-MSH are skin pigmentation *via* MC1-R in melanocytes and regulation of energy homeostasis *via* MC4-R in the brain.^10,11^ Owing to these actions, the principal sites of POMC expression and its processing into biologically active α-MSH lie within the central nervous system (hypothalamus and pituitary gland) and in the skin, but there is some evidence demonstrating that α-MSH is also produced in other peripheral tissues.^12^ Post-translational processing of POMC into mature α-MSH necessitates well-coordinated actions of several enzymes including carboxypeptidase E (CPE) and α-amidating monooxygenase (PAM), which co-localize in POMC-expressing cells.^7^ It is generally thought that α-MSH is released into circulation by the pituitary gland, while in other tissues, α-MSH acts in an autocrine or paracrine fashion. Interestingly, early studies have discovered that POMC mRNA and α-MSH production also occur in the rat heart.^13,14^ As a relevant clinical observation, plasma α-MSH level was found to be elevated in patients suffering from hypertrophic or dilated cardiomyopathy.^15^ However, the role of α-MSH and its possible target receptors in the heart are completely unknown.

Given the expression of α-MSH in the rat heart and increased plasma levels in heart failure patients, we hypothesized that its production in the heart is sensitive to pressure overload and modulated during the development of heart failure. We found that α-MSH production is significantly reduced in the failing mouse heart and that administration of a stable α-MSH analogue protects the mice against pressure overload-induced cardiac hypertrophy and heart failure. Experimental data from a series of *in vitro* and *in vivo* experiments show that α-MSH acts as an antihypertrophic regulator by interacting with MC5-R in cardiac myocytes.

## Methods

### Mice and Study Design

Mice were housed in groups on a 12h light/dark cycle with free access to food (# 2916C, Teklad Global diet, Envigo) and tap water. The experiments were approved by the national Animal Experiment Board in Finland (License number: ESAVI/6280/04.10.07/2016 and ESAVI/1260/2020) and conducted in accordance with the Directive 2010/63/EU of the European Parliament on the protection of animals used for scientific purposes and with the institutional and national guidelines for the care and use of laboratory animals.

For nonselective activation of the melanocortin system *in vivo*, 8-week-old male C57BL/6J mice were subjected to transverse aortic constriction (TAC) as described below. Mice were allowed to recover for 2 weeks from the surgery and were thereafter randomly assigned to receive daily i.p. injections of either vehicle (PBS) or the α-MSH analogue melanotan-II (0.3 mg/kg/day, Tocris, # 2566).^16^ Sham-operated mice received vehicle according to the same treatment scheme. Mice were sacrificed 8 weeks after the TAC operation.

To generate inducible cardiomyocyte-specific MC5-R knockout (Mc5r-cKO) mice, MC5-R floxed mice (Mc5r^fl/fl^, GemPharmatec, strain # T00591) were intercrossed with tamoxifen-inducible Myh6-MerCreMer transgenic mice (Myh6-MCM, the Jackson Laboratory, strain # 005657).^17^

To study the therapeutic benefits of MC5-R activation in a heart failure model, 8-week-old male C57BL/6N were subjected to TAC and after a one-week recovery period, randomly assigned to receive i.p. injections of either vehicle (PBS) or the selective MC5-R agonist PG-901 (0.005 or 0.5 mg/kg/day). Mice were sacrificed 5 weeks after the TAC operation.

Further details on study design and experimental procedures are given in the Supplemental Material.

### Cardiac Hypertrophy Models and Echocardiography

To induce hemodynamic pressure overload, mice were subjected to TAC as previously described.^18^ As another model of cardiac hypertrophy, mice were subjected to subcutaneous infusion of angiotensin II (1.4 mg/kg/day) for 2 or 4 weeks using osmotic minipumps (Alzet, Model 1004).^18^ For the surgical operations, mice were anesthetized with ketamine (110 mg/kg, i.p.) and xylazine (15 mg/kg, i.p.), and buprenorphine (0.05 mg/kg, s.c., 2x/day for 3 days) and carprofen (5 mg/kg, s.c., 1x/day for 3 days) were given for peri- and post-operative analgesia. Cardiac structure and function were assessed by transthoracic echocardiography (Vevo 2100, Visual Sonics Inc., Toronto, Canada) before the start of drug administration and at the end of the experiment under isoflurane anesthesia (4 % for induction and 2 % for maintenance). Echocardiography measurements of cardiac structure and function prior to drug administration are reported in **Supplementary Tables I-II**

### Single-Cell RNA Sequencing Analysis

Single-cell RNA-sequencing data deposited in the Gene Expression Omnibus (GSE120064) was used to study *Pomc*-, *Cpe*- and *Pam*-expressing cell types in the mouse heart. All quality control passed cells (11 492) were analyzed using Seurat (version 4.0.3) with R (version 4.2.0).^19^ Raw gene counts were first normalized using LogNormalize method and then scaled using a scale factor of 10 000. An unsupervised dimensional reduction method, UMAP (Uniform Manifold Approximation and Projection) was used to generate clustering results using first 26 principal components. The same cell type annotation as reported by the original authors was used in the analysis.^20^

### Cell Culture and Treatments

Rat heart myoblast H9c2(2-1) cells (ATCC®, CRL-1446™), neonatal mouse ventricular cardiac myocytes (NMCM) and human induced pluripotent stem cell-derived cardiomyocytes (hiPSC-CM)^21,22^ were used to study the effects of α-MSH (abcam, # ab120189) and selective MC-R agonists *in vitro*. LD211 was used as a selective agonist for MC1-R (compound 28 in the original publication),^23^ [D-Trp8]-γ-MSH as a selective MC3-R agonist,^24^ THIQ as a selective MC4-R agonist (Cayman Chemical, USA, # 312637-48-2)^25^ and PG-901 as a selective MC5-R agonist.^26^ PG-20N was used as a selective MC5-R antagonist.^27^ LD211, [D-Trp8]-γ-MSH, PG-901 and PG-20N were synthetized and provided by professor Minying Cai.

### Human Cardiomyopathy Samples

Human LV samples were obtained from dilated (n=15) and ischemic (n=8) cardiomyopathy patients undergoing cardiac transplantation in Helsinki University Hospital between 2014-2019. The patient characteristics have been previously reported.^28^ Control samples (n=13) were from victims of traffic accidents with no history or evidence of cardiovascular diseases at autopsy. The study was approved by the Ethics Committee of Helsinki and Uusimaa Hospital District and conducted according to the declaration of Helsinki, and the study subjects gave informed consent.

### RNA Isolation, cDNA Synthesis and Quantitative RT-PCR

H9c2, NMCMs and hiPSC-CMs were collected into QIAzol Lysis Reagent or Trizol Reagent (Invitrogen) and total RNA was extracted using Direct-zol RNA Miniprep and Microprep (Zymo Research, CA, USA), respectively. RNA was reverse-transcribed to cDNA (PrimeScript RT reagent kit, Takara Clontech) and quantitative real-time polymerase chain reaction (RT-PCR) was performed with SYBR Green protocols (Kapa Biosystems, MA, USA) and a real-time PCR detection system (Applied Biosystems 7300 Real-Time PCR system).^29,30^ Target gene expression was normalized to a housekeeping gene (ribosomal protein S18; RPS18, glyceraldehyde-3-phosphate dehydrogenase; GAPDH or β-actin; ACTB) using the comparative ΔCt method and results are presented as relative transcript levels (2^-ΔΔCt^). Primer sequences are presented in **Supplementary Tables III-V**.

### Statistics

Statistical analyses were performed with GraphPad Prism 8 software (La Jolla, CA, USA). Statistical significance between the experimental groups was determined by unpaired Student’s t-test, one-way ANOVA followed by Dunnett *post hoc* tests or two-way ANOVA followed by Bonferroni *post hoc* tests. Quantitative PCR data from hiPSC-CMs was analyzed using ΔCt-values and randomized block ANOVA (individual experiments and treatment as factors) as previously described.^31^ Pearson correlation coefficients were calculated for gene associations. The D’Agostino and Pearson omnibus normality test method was employed to test the normality of the data. Possible outliers in the data sets were identified using the regression and outlier removal (ROUT) method at Q-level of 1 %. Data are expressed as mean ± standard error of the mean (SEM). Results were considered significant for P<0.05.

## Results

### α-MSH is Expressed in the Heart and Protects against Pressure Overload-Induced Cardiac Hypertrophy in Mice

We first aimed to investigate whether α-MSH is expressed in the mouse heart. Using an ELISA assay, we found that cardiac α-MSH concentration is comparable to its level in the hypothalamus (**Figure 1A**) that is considered as a rich source of α-MSH. α-MSH expression appeared to be also higher in the ventricles compared to the atria (**Figure 1A**). In contrast, α-MSH concentration in other reference tissues such as the spleen and skeletal muscle was considerably lower (**Figure 1A**). Cardiac α-MSH expression was also confirmed by immunohistochemistry, which revealed a small subset of cardiac cells that were positive for α-MSH (**Figure 1B**). Using single-cell RNA sequencing data from TAC-operated mice,^20^ we performed clustering analysis to identify *Pomc*-, *Cpe*- and *Pam*-expressing cells in major cell types of the heart (**Figure 1C, Supplementary Figure I**). Although *Pomc^+^* -cells were identified among all cell clusters, the majority of the triple-positive *Pomc*^+^ *Cpe*^+^ *Pam*^+^ -cells, which are likely to produce POMC into mature α-MSH, were cardiomyocytes (**Figure 1D**). Next, to test the hypothesis that α-MSH production is triggered by pressure overload, we analyzed changes in cardiac *Pomc*, *Cpe* and *Pam* expression at different stages of cardiac hypertrophy. The relative number of *Pomc^+^* -cells increased consistently in all major cell types during the early stage of cardiac hypertrophy (2 weeks after TAC) and then declined during the development of heart failure (5-8 weeks after TAC) (**Figure 1E**). Likewise, in terms of changes in *Pomc*^+^ *Cpe*^+^ *Pam*^+^ -cardiomyocyte count, there was a biphasic response with an initial elevation and then a decline in the failing heart (**Figure 1F**). In good agreement with these findings, α-MSH concentration in the LV was reduced at the late stage of hypertrophy (5 weeks after TAC) (**Figure 1G**), an effect occurring without a change in plasma α-MSH concentration (**Supplementary Figure I**). The TAC-induced reduction of ventricular α-MSH expression was also confirmed by Western blotting (**Supplementary Figure I**).

**Figure 1.**
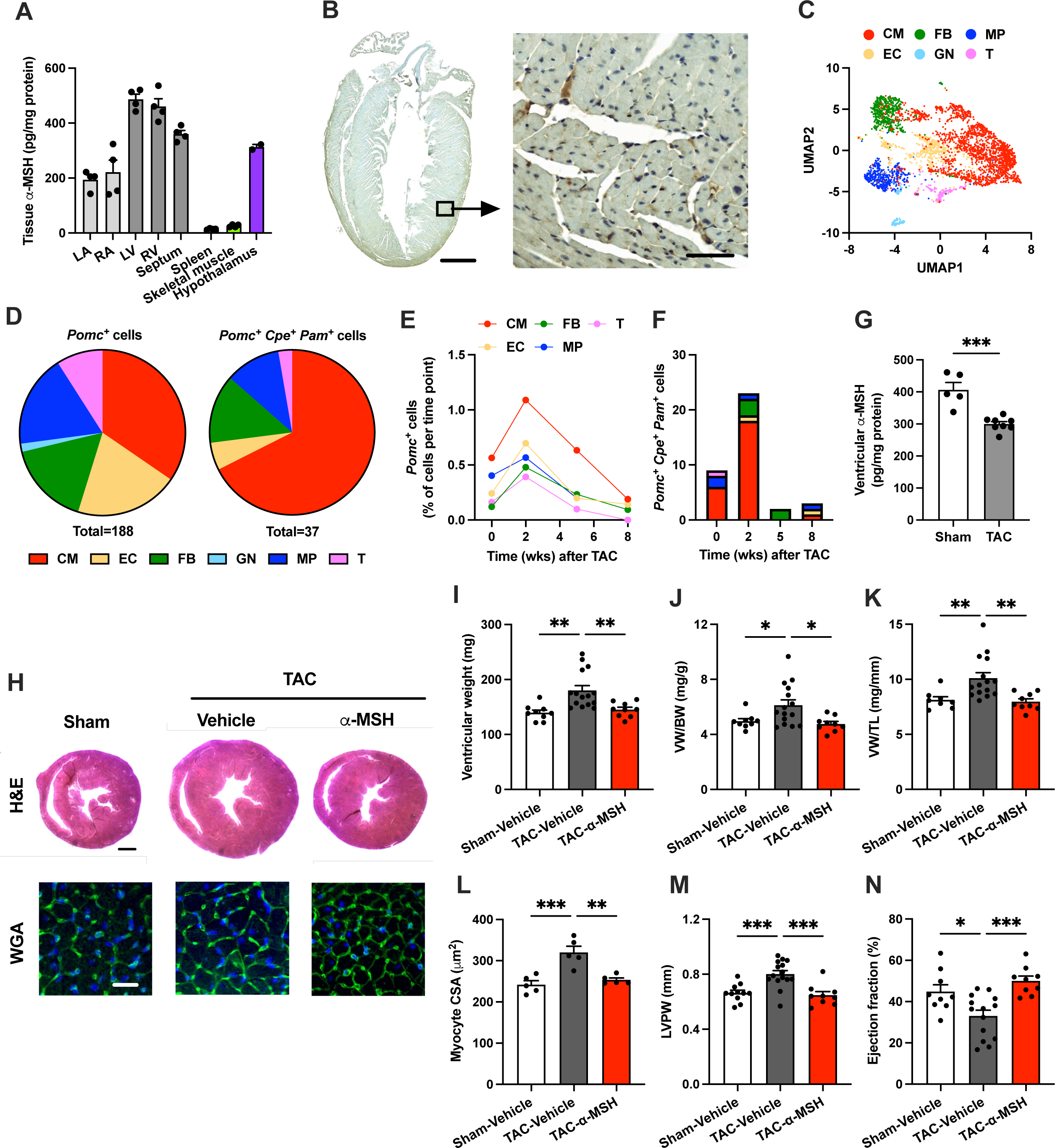
α-Melanocyte-stimulating hormone (α-MSH) is produced in the heart and protects against pathological cardiac hypertrophy. **A**, α-MSH concentration (pg/mg protein) in the mouse left atrium (LA), right atrium (RA), left ventricle (LV), right ventricle (RV), septum, spleen, skeletal muscle and medial basal hypothalamus. **B**, Immunostaining of α-MSH in longitudinal heart section of C57Bl/6J mouse. Scale bar, 1 mm (left) and 50 μm (right). **C**, Uniform Manifold Approximation and Projection (UMAP) showing 11 492 single cells isolated from C57Bl mice at different stages of cardiac hypertrophy (0-8 weeks after TAC). Cell types were determined according to the expression of known markers. CM indicates cardiomyocyte; EC, endothelial cell; FB, fibroblast; GN, granulocyte; MP, macrophage; and T, T cell. **D**, Pie charts showing the relative distribution of Pomc-positive (*Pomc*^+^) and triple-positive (*Pomc*^+^ *Cpe*^+^ *Pam*^+^) cells in each cell type. Cells were pooled from all time points (0-8 weeks after TAC). **E**, Changes in the relative amount of *Pomc*^+^ -cells in each cell type as a function of time after transverse aortic constriction (TAC) surgery. *Pomc*^+^ -cells are expressed as percentage of total number of sequenced cells at each time point. **F**, Changes in the number of *Pomc*^+^ *Cpe*^+^ *Pam*^+^ -cells in each cell type as a function of time after TAC surgery. **G**, α-MSH concentration in the LV of sham- and TAC-operated mice 5 weeks after the surgery. *** *P*<0.001 by Student’s t test. **H**, Exemplary hematoxylin and eosin (H&E)-stained cross-sections of the heart showing the gross morphology of sham- and TAC-operated mice treated with either vehicle or α-MSH analogue (melanotan II; MT-II). Scale bar, 1 mm. Lower panel of images show exemplary cross-sections of the heart stained with wheat germ agglutinin (WGA) to demarcate cell boundaries. Scale bar, 25 μm. **I** through **K**, Ventricular weight, ventricular weight to tibia length ratio (VW/TL) and ventricular weight to body weight ratio (VW/BW) in the indicated groups. **L**, Quantification of cross-sectional area of ventricular cardiomyocytes. **M** and **N**, Left ventricular posterior wall thickness (LVPW) and ejection fraction analyzed by echocardiography at the end of the experiment. Data are mean ± SEM, each dot represents individual mouse. * *P*<0.05, ** *P*<0.01 and *** *P*<0.001 for the indicated comparisons by 1-way ANOVA and Dunnett *post hoc* tests.

The declining level of α-MSH in the failing heart raised a question whether α-MSH could be protective against pathological cardiac hypertrophy. To investigate the therapeutic potential of α-MSH, we subjected C57BL/6J mice to TAC surgery and randomly assigned the mice to receive daily injections of either vehicle or a stable analogue of α-MSH (melanotan-II; MT-II). After 8 weeks of TAC, α-MSH-treated mice showed attenuated LV hypertrophy compared to vehicle-treated TAC mice as evidenced by reduction in ventricular weight, ventricular weight-to-tibia length ratio and ventricular weight-to-body weight ratio (**Figure 1H-K**). Histological examination also revealed reduced cardiomyocyte size in α-MSH-treated mice (**Figure 1H** and **1L**). As determined by echocardiography, α-MSH treatment prevented TAC-induced thickening of LV posterior wall (**Figure 1M**) and deterioration of LV ejection fraction (**Figure 1N**). Lastly, gene expression analysis demonstrated that the molecular fingerprint of pathological hypertrophy was partly reversed by α-MSH with a significant reduction in TAC-induced expression of fibrosis-related genes including *Col1a2* (collagen type I, alpha 2) and *Mmp2* (matrix metalloproteinase-2) (**Supplementary Figure II**). Taken together, these data indicate that local α-MSH production in the heart is responsive to pressure overload and that α-MSH acts as an antihypertrophic regulator.

### Cardiomyocytes Are Responsive to Treatment with α-MSH

To investigate whether functional MC-Rs exist in cardiomyocytes, we performed experiments with H9c2 cells and NMCMs. Cells were treated with α-MSH for 5, 15, 30 or 60 minutes and assayed for intracellular cAMP levels since most MC-Rs are known to be coupled to G_s_ proteins and cAMP signaling. Unexpectedly, α-MSH caused a reduction in cAMP level at 5 min time point in H9c2 cells and NMCMs (**Figure 2A**), indicating Gi-dependent coupling of the α-MSH response. Screening of other potential downstream targets of melanocortin signaling revealed a marked reduction in JNK phosphorylation after α-MSH treatment (**Figure 2B**), while no effect was observed on phosphorylation of ERK1/2 (**Figure 2C**) or intracellular Ca^2+^ responses (**Supplementary Figure III**). In terms of concentration-responsiveness, α-MSH reduced intracellular cAMP level under baseline conditions (**Figure 2D**) as well as in cells stimulated with the adenylyl cyclase activator forskolin (**Figure 2E**) with effect peaking in the subnanomolar range of concentrations (< 1 nM or log −9 M). JNK phosphorylation was also reduced at subnanomolar concentrations of α-MSH (**Figure 2F**).

**Figure 2.**
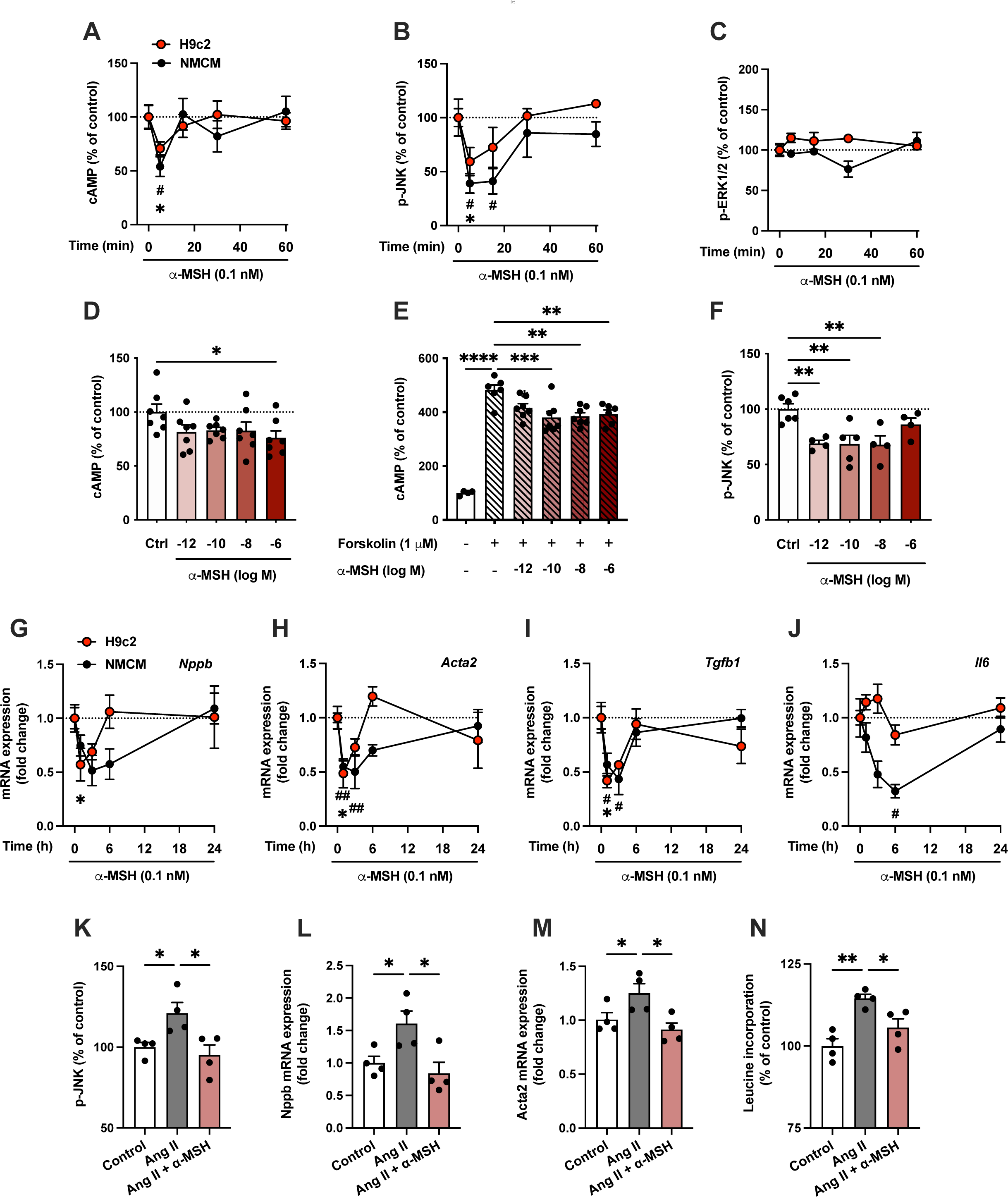
α-MSH reduces the levels of cAMP and phosphorylated JNK in cultured cardiomyocytes. **A**, Quantification of intracellular cAMP levels in H9c2 cells and neonatal mouse ventricular cardiac myocytes (NMCMs) treated with α-MSH (0.1 nM) for 5, 15, 30 or 60 minutes. Data is expressed as percentage of control (Ctrl, 0 min). **B** and **C**, Quantification of phosphorylated JNK and ERK1/2 by ELISA assays in H9c2 cells and NMCMs treated with α-MSH (0.1 nM) for 5, 15, 30 or 60 minutes. * *P*<0.05 versus Control (0 min) in H9c2 cells, # *P*<0.05 versus Control (0 min) in NMCMs by 1-way ANOVA and Dunnett *post hoc* tests. **D** and **E,** Quantification of intracellular cAMP levels in H9c2 cells treated with different concentrations of α-MSH for 30 minutes in the absence or presence of forskolin (1 μM). **F**, Quantification of phosphorylated JNK using ELISA assay in H9c2 cells treated with different concentrations of α-MSH for 30 minutes. * *P*<0.05, ** *P*<0.01, *** *P*<0.001 and **** *P*<0.0001 for the indicated comparisons by 1-way ANOVA and Dunnett *post hoc* tests. **G** through **J**, Quantitative real-time PCR (qPCR) analysis of *Nppb* (B-type natriuretic peptide), *Acta2* (alpha-smooth muscle actin), *Tgfb1* (transforming growth factor beta 1) and *Il6* (interleukin 6) in H9c2 cells and NMCMs treated with α-MSH (0.1 nM) for 1, 3, 6 or 24 hours. * *P*<0.05 versus Control (0 h) in H9c2 cells, # *P*<0.05 and ## *P*<0.01 versus Control (0 h) in NMCMs by 1-way ANOVA and Dunnett *post hoc* tests. **K** through **M**, Quantification of phosphorylated JNK by ELISA assay and qPCR analysis of *Nppb* and *Acta2* expression in H9c2 cells treated with angiotensin II (Ang II, 0.1 μM) in the absence or presence of α-MSH (0.1 nM). **N**, [^3^H]-Leucine incorporation assay in H9c2 cells treated with Ang II (0.1 μM) for 24 hours in the absence or presence of α-MSH (0.1 nM). * *P*<0.05 and ** *P*<0.01 for the indicated comparisons by 1-way ANOVA and Dunnett *post hoc* tests. Data are mean ± SEM, n=4-7 per group in each graph.

Finally, to investigate whether the activation of these signaling cascades leads to changes in gene expression, we performed qPCR analysis for H9c2 cells and NMCMs that were treated with α-MSH for 1, 3, 6 or 24 hours. We observed downregulation of the hypertrophy related gene *Nppb* (B-type natriuretic peptide, **Figure H**) and fibrosis-related genes including *Acta2* (alpha-smooth muscle actin, **Figure 2I**), *Tgfb1* (transforming growth factor beta 1, **Figure 2J**), *Col3a1* (collagen type III, alpha 1, **Supplementary Figure III**) and *Fn1* (fibronectin, **Supplementary Figure III**). In addition, the pro-inflammatory gene *Il6* (interleukin 6) was downregulated in NMCMs but not in H9c2 cells (**Figure 2K**). In general, the changes in gene expression were short-lasting and vanished within 24 hours, which probably reflects the short half-life of α-MSH.

To investigate the effects of α-MSH in hypertrophic context, we treated H9c2 cells with angiotensin II (Ang II) to promote cellular hypertrophy and used leucine incorporation assay as a measure of protein synthesis rate. α-MSH effectively prevented Ang II-induced increase in leucine incorporation (**Figure 2K**). α-MSH also reversed the increase in JNK phosphorylation as well as in *Nppb* and *Acta2* expression in Ang II-treated cells (**Figure 2L-N**). Collectively, the results demonstrate that functional MC-Rs exist in cardiomyocytes and suggest that the antihypertrophic effect observed in TAC-operated mice is dependent on direct actions of α-MSH on cardiomyocytes.

### MC5-R Is Expressed in Mouse Cardiomyocytes and Downregulated During Development of Heart Failure

To investigate which MC-R subtype mediates the effects of α-MSH in cardiomyocytes, we used subtype selective MC-R agonists and screened their effects on gene expression by qPCR. We found that the MC5-R selective agonist PG-901 was the only compound that clearly mimicked the actions of α-MSH (**Figure 3A-D, Supplementary Figure IV**). Agonism at MC5-R downregulated *Tgfb1, Col3a1* and *Fn1,* and tended to also reduce *Acta1* (actin, alpha skeletal muscle) and *Acta2* mRNA levels (*P*=0.08 and 0.09, respectively) (**Figure 3A-D, Supplementary Figure IV**). We then sought to determine whether the α-MSH-induced effects are dependent on MC5-R activation in cardiomyocytes. Addition of the selective MC5-R antagonist PG-20N completely reversed the effect of α-MSH on p-JNK level and gene expression changes (**Figure 3E-H**), further supporting the role of MC5-R as a mediator of the α-MSH-induced effects.

**Figure 3.**
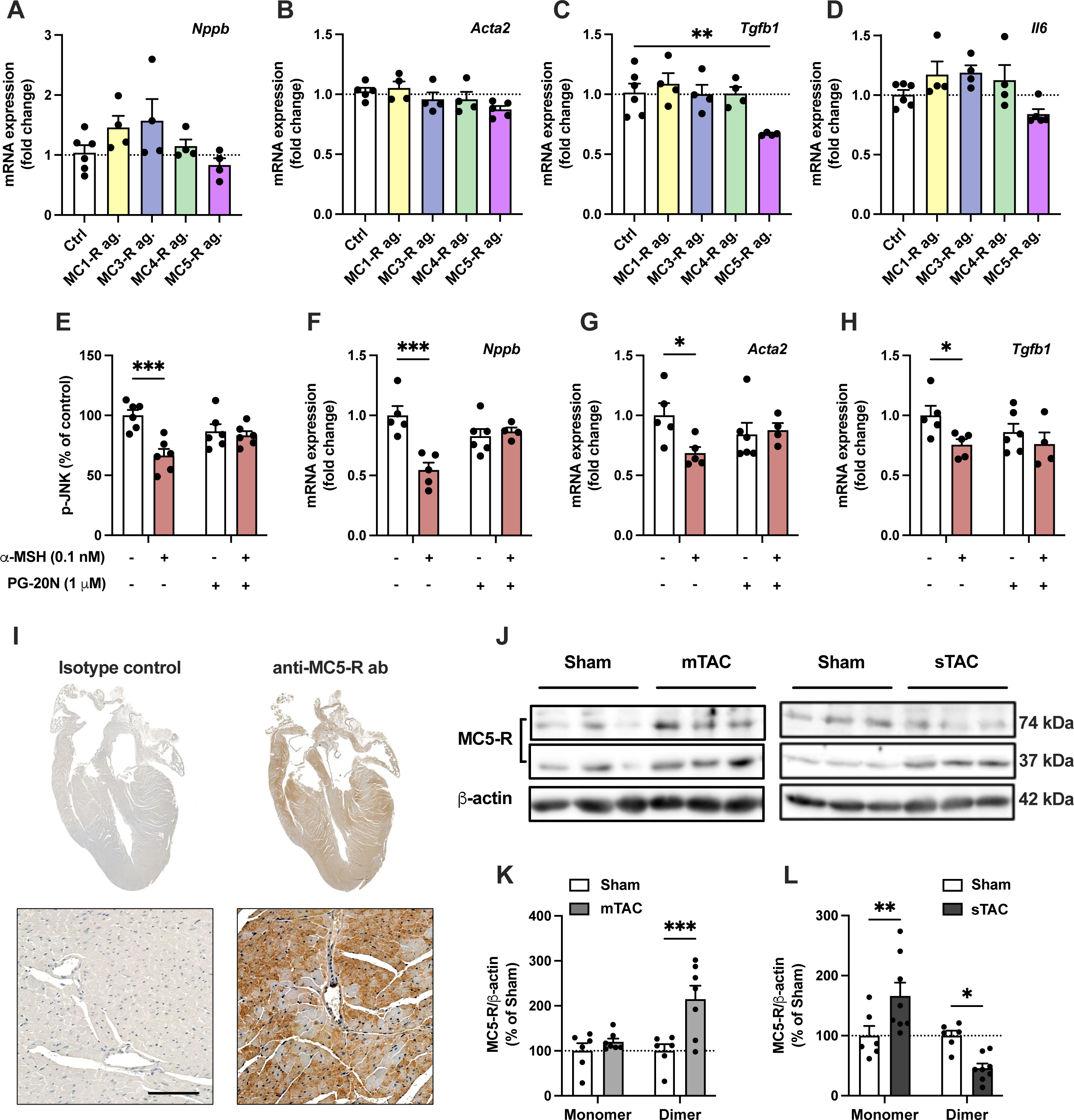
MC5-R is expressed in the mouse heart and its level is regulated by pressure overload. **A** through **D**, Quantitative real-time PCR (qPCR) analysis of *Nppb* (B-type natriuretic peptide), *Acta2* (alpha-smooth muscle actin), *Tgfb1* (transforming growth factor beta 1) and *Il6* (interleukin 6) mRNA expression in H9c2 cells treated with subtype selective MC-R agonists for 3 hours. n=4-6 per group in each graph. **E** through **H**, Quantification of phosphorylated JNK by ELISA assay and qPCR analysis of *Nppb*, *Acta2* and *Tgfb1* expression in H9c2 cells treated with α-MSH (0.1 nM) for 1 hour in the absence or presence of the selective MC5-R antagonist PG-20N (1 μM). n=4-6 per group in each graph. **I**, Immunostaining of MC5-R in longitudinal heart section of C57Bl/6J mouse. In control section, anti MC5-R antibody was replaced by purified normal rabbit IgG (isotype control). Scale bar, 100 μm. **J**, Representative Western blots showing the monomer (37 kDa) and dimer (74 kDa) forms of MC5-R and β-actin (loading control) in the left ventricle (LV) after sham or TAC surgery. mTAC indicates mild TAC; sTAC, severe TAC. **K** and **L**, Quantification of MC5-R monomer and dimer forms (normalized against β-actin) in mTAC and sTAC LV samples. n=6-8 mice per group in each graph, each dot represents individual mouse. * P<0.05, ** *P*<0.01, *** *P*<0.001 for the indicated comparisons by 1-way ANOVA and Dunnett *post hoc* tests (**C**) or by 2-way ANOVA and Bonferroni *post hoc* tests (**E**-**L**).

We next aimed to explore whether MC5-R is expressed in the mouse heart. Immunohistochemical staining revealed that MC5-R is uniformly present in the heart (**Figure 3I**). We also studied whether MC5-R protein abundance is changed in pressure-overloaded LV samples. For this purpose, we preselected samples from TAC-operated mice that displayed mild hypertrophy with normal LV systolic function (EF −3 % vs sham, P=0.23) and another set of mice which had more advanced hypertrophy and significant LV dysfunction (EF −25 % vs sham, P<0.001). Western blotting analysis revealed that in mildly hypertrophied LV samples (referred to as mTAC), MC5-R dimer, which is considered as functionally more active form,^32^ was significantly upregulated, while no change was found in the expression of MC5-R monomer (**Figure 3J** and **3K**). In contrast, MC5-R dimer was markedly reduced in severely hypertrophied LV samples (referred to as sTAC) and it was accompanied by a slight increase in the amount of MC5-R monomer (**Figure 3J** and **3L).** Cardiac hypertrophy induced by Ang II infusion (for 2 or 4 weeks) did not however affect MC5-R protein level in the LV (**Supplemental Figure IV**).

### MC5-R Acts as an Anti-Hypertrophic Regulator

Since MC5-R appeared to be the target receptor for α-MSH in the heart, we turned our attention more closely to the MC5-R agonist PG-901 and investigated its actions on intracellular signaling cascades in cultured cardiomyocytes. In line with the effects evoked by α-MSH, PG-901 inhibited JNK pathway as evidenced by reduction in the phosphorylated form of JNK (**Figure 4A**). No significant changes were observed in the phosphorylation of ERK1/2 (**Figure 4B)** or intracellular Ca2+ levels after PG-901 treatment (**Supplementary Figure V**). In terms of JNK inhibition, the effect was most potent at 0.1 nM (log −10 M) concentration of PG-901 in H9c2 cells (**Figure 4C**). Similarly to α-MSH, PG-901 reduced intracellular cAMP level under baseline conditions (**Figure 4D**) and to some extent also in forskolin-stimulated cells (**Supplementary Figure V**). However, under a stronger and more physiological stimulus induced by the β-adrenoceptor agonist isoprenaline, PG-901 had no effect on cAMP level (**Supplementary Figure V**), suggesting a rather weak coupling of MC5-R to Gi protein. In terms of gene expression, PG-901 downregulated *Acta1*, *Col3a1* and *Tgfb1* in H9c2 cells with a concentration profile similar to JNK inhibition (**Supplementary Figure V**). Other genes such as *Fn1*, *Col1a1*, *Col1a2*, *Ctgf* (connective tissue growth factor) and *Mmp2* were downregulated most significantly at the lowest concentration of PG-901 (1 pM or log −12 M) (**Supplementary Figure V**).

**Figure 4.**
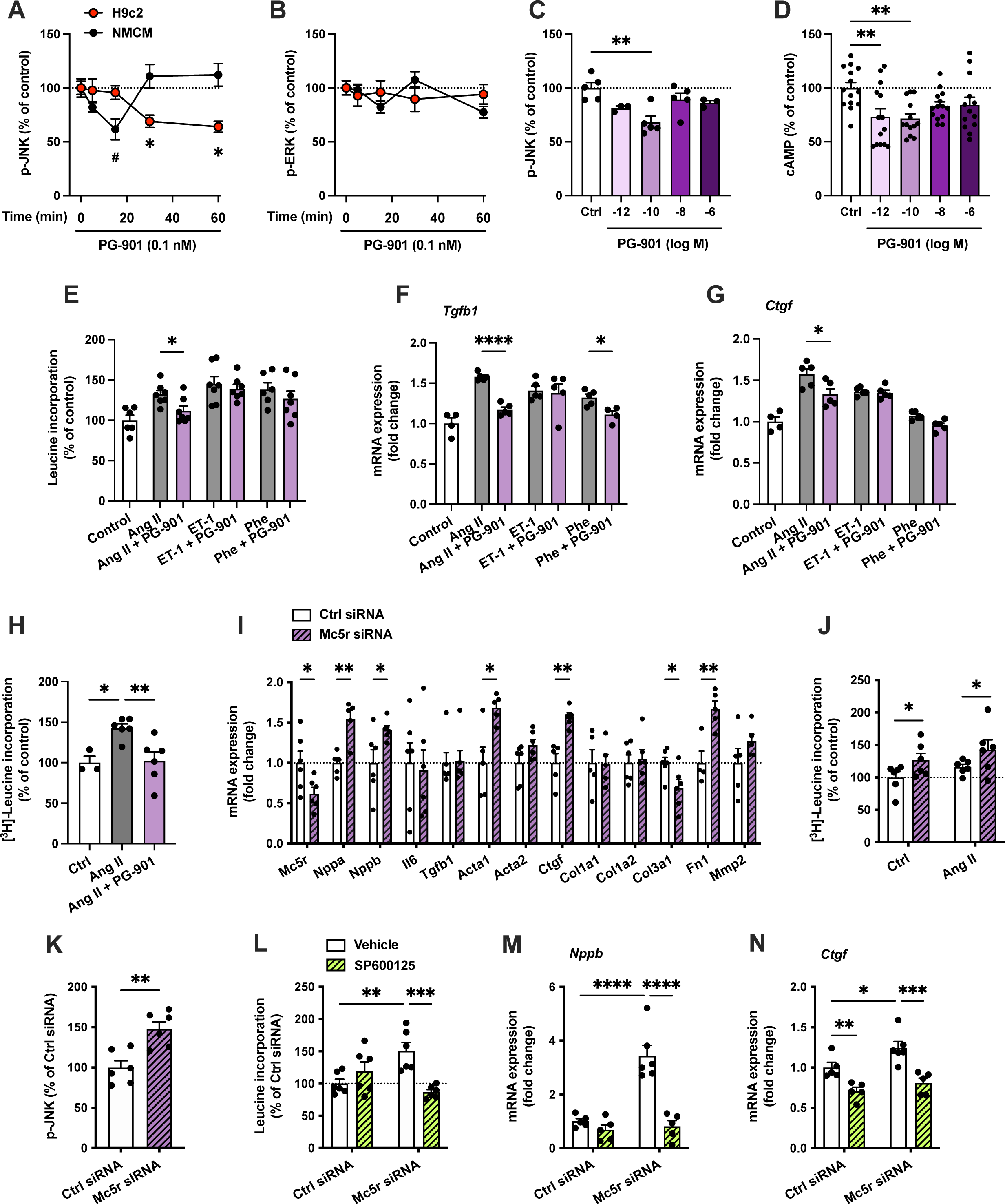
MC5-R activation with PG-901 mimics the actions of α-MSH in cultured cardiomyocytes. **A** and **B**, Quantification of phosphorylated JNK and ERK by ELISA assays in H9c2 cells and NMCMs treated with PG-901 (0.1 nM) for 5, 15, 30 or 60 minutes. Data is expressed as percentage of control (0 min). * *P*<0.05 versus Control (0 min) in H9c2 cells, # *P*<0.05 versus Control (0 min) in NMCMs. n=4-6 per time point. **C**, Quantification of phosphorylated JNK using ELISA assay in H9c2 cells treated with different concentrations of PG-901 for 60 minutes. n=4-6 per group. **D,** Quantification of intracellular cAMP levels in H9c2 cells treated with different concentrations of PG-901 for 30 minutes. n=13-14 per group. **E**, [^3^H]-Leucine incorporation assay in H9c2 cells treated with angiotensin II (Ang II, 0.1 μM), endothelin 1 (ET-1, 0.1 μM) or phenylephrine (Phe, 0.1 mM) for 24 hours in the absence or presence of PG-901 (0.1 nM). n=5-7 per group. **F** and **G**, Quantitative real-time PCR (qPCR) analysis of *Tgfb1* and *Ctgf* mRNA expression in H9c2 cells treated with Ang II, ET-1 or Phe for 3 hours in the absence or presence of PG-901 (0.1 nM). n=4-5 per group. **H**, [^3^H]-Leucine incorporation assay in NMCMs treated with Ang II for 24 hours in the absence or presence of PG-901 (0.1 nM). n=3-6 per group. **I**, qPCR analysis of the indicated genes in NMCMs treated with control siRNA or *Mc5r* targeting siRNA for 24 hours. n=4-6 per group**. J**, [^3^H]-Leucine incorporation in NMCMs treated with control siRNA or *Mc5r* targeting siRNA for 24 hours. **K**, Quantification of phosphorylated JNK in H9c2 cells treated with control siRNA or *Mc5r* targeting siRNA for 24 hours. **L**, [^3^H]-Leucine incorporation in H9c2 cells treated with or without the JNK inhibitor SP600125 (10 μM) for 30 minutes followed by transfection with control siRNA or *Mc5r* targeting siRNA for 24 hours. **M** and **N** qPCR analysis of *Nppb* and *Ctgf* mRNA expression in H9c2 cells treated with or without the JNK inhibitor SP600125 (10 μM) for 30 minutes followed by transfection with control siRNA or *Mc5r* targeting siRNA for 24 hours. n=6 per group. Data are mean ± SEM, *P*<0.05, ** *P*<0.01 and **** *P*<0.0001 for the indicated comparisons by Student’s t-test (**I** and **K**), 1-way ANOVA and Dunnett *post hoc* tests (**E-H** and **J**) or 2-way and Bonferroni *post hoc* tests (**L**-**N**).

Further mechanistic experiments using H9c2 cells revealed that inhibition of Gi signaling with pertussis toxin (PTX) induced a further reduction in the amount of phosphorylated JNK and downregulation of *Nppb* and *Acta2*. Although PTX appeared to block the effect of PG-901 on JNK phosphorylation, it did not abrogate the gene expression changes evoked by MC5-R activation (**Supplementary Figure VI**), indicating that signaling *via* Gi pathway does not mediate the downstream effects of MC5-R activation. Since MC5-R can simultaneously signal through both Gs- and Gi-dependent pathways,^33^ we also tested whether the effects of PG-901 could be reversed by blocking the cAMP-PKA axis. The PKA inhibitor H89 or a cAMP analog (cAMPS-Rp) that antagonizes cAMP-induced activation of PKA did not abolish the PG-901 evoked reduction of p-JNK, but caused a further lowering of p-JNK level (**Supplementary Figure VII**). Gene expression changes associated with MC5-R activation were not either affected by inhibition of PKA (**Supplementary Figure VII**). By contrast, the JNK activator anisomycin completely reversed the effect of PG-901 on fibrosis-associated genes such as *Acta2*, *Tgfb1*, *Col1a2*, *Ctgf* and *Fn1* (**Supplementary Figure VIII**). Taken together, these results demonstrate that the MC5-R-evoked changes in gene expression occur in a cAMP-independent manner but are at least partly dependent on the JNK pathway.

Finally, to investigate whether MC5-R has a physiological significance in the regulation of cardiomyocyte growth, we used leucine incorporation assay and treated H9c2 cells with PG-901 in combination with different hypertrophic stimuli including Ang II, ET-1 and the α-adrenoceptor agonist phenylephrine (Phe). We found that PG-901 effectively prevented Ang II-induced increase in leucine incorporation (**Figure 4E**), which is largely dependent on JNK activation.^34^ In contrast, no significant effect was observed on ET-1- or Phe-induced hypertrophic response (**Figure 4E**), which has been shown to be more dependent on p38 phosphorylation.^35^ Likewise, in terms of gene expression, PG-901 blunted the induction of fibrotic genes such as *Tgfb1* and *Ctgf* most notably in Ang II-stimulated cells (**Figure 4F, 4G** and **Supplementary Figure IX**). The inhibitory effect of PG-901 on leucine incorporation was also confirmed in Ang II-stimulated NMCMs (**Figure 4H**). We next investigated whether MC5-R silencing by siRNA causes an opposite phenotype to that seen after MC5-R activation. Indeed, MC5-R knockdown was associated with upregulation of *Nppa* (atrial natriuretic peptide), *Nppb*, *Acta1, Ctgf* and *Fn1* (**Figure 4I**) as well as with enhanced leucine incorporation (**Figure 4J**) and JNK phosphorylation (**Figure 4K**). To test the dependency of the observed phenotype on JNK signaling, H9c2 cells were treated with the selective JNK inhibitor SP600125 prior to transfection with the *Mc5r* targeting siRNA. We observed that JNK inhibition with SP600125 completely abolished the increase in leucine incorporation and upregulation of *Nppb* and *Ctgf* in *Mc5r*-silenced cells (**Figure 4L-N**), further consolidating the link between MC5-R signaling, JNK pathway and cardiomyocyte hypertrophy.

### Human Cardiomyocytes Express Functional MC5-R

The findings implicating an antihypertrophic role for MC5-R in mouse cardiomyocytes prompted us to investigate whether human cardiomyocytes also express MC5-R. We first quantified the mRNA levels of *MC5R* in the LV samples from control subject and from patients with end-stage dilated (DCM) or ischemic cardiomyopathy (ICM). *MC5R* was significantly upregulated in the LV of DCM and ICM patients compared to controls (**Figure 5A**). Likewise, *POMC* expression was increased in the diseased human hearts, particularly in the ICM patients (**Figure 5B**). We also found that *MC5R* and *POMC* mRNA levels correlated positively and highly significantly in the DCM and ICM samples (**Figure 5C**). We next studied whether *MC5R* and *POMC* expression are regulated by different hypertrophic stimuli in hiPSC-CMs. First, hiPSC-CMs were exposed to mechanical stretch for 24 or 48 hours,^36^ which led to a distinct gene expression pattern of the natriuretic peptides with a delayed upregulation of *NPPA* (after 48 hours) and a more rapid induction of *NPPB* (after 24 hours) (**Supplementary Figure X**). Intriguingly, mechanical stretching of hiPSC-CMs downregulated *MC5R* expression but this occurred only after 48 hours of stretch (**Figure 5D**). In contrast, no change was observed in POMC expression (**Figure 5E**). As another model of cardiomyocyte hypertrophy, hiPSC-CMs were treated with ET-1 for 24 hours,^37^ which led to stronger induction of *NPPA* and *NPPB* expression compared to mechanical stretching (**Supplementary Figure X**). ET-1 treatment also clearly downregulated *MC5R,* while *POMC* expression was unaffected (**Figure 5F and 5G**), corroborating the finding from mechanical load-induced hypertrophy of hiPSC-CMs. Although no change was observed in *POMC* expression, *MC5R* expression positively correlated with *POMC* mRNA levels in the ET-1-treated hiPSC-CM samples (**Supplementary Figure X**), suggesting transcriptional co-regulation of these genes in cultured cardiomyocytes as well as in the human heart.

**Figure 5.**
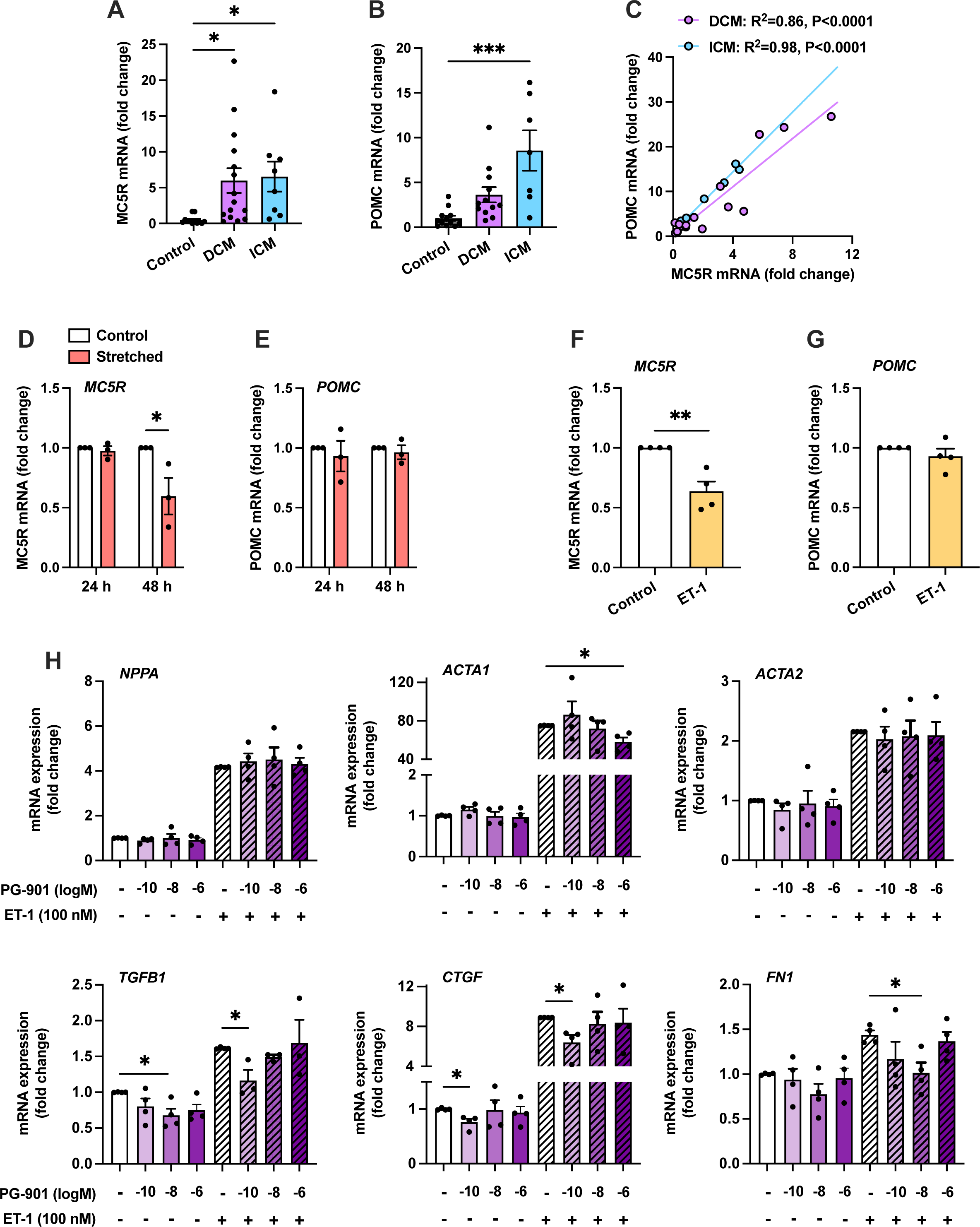
*MC5R* expression in human LV samples from cardiomyopathy patients and in human induced pluripotent stem cell-derived cardiomyocytes. **A** and **B**, Quantitative PCR analysis of *MC5R* and *POMC* mRNA expression in left ventricular (LV) samples from healthy control subjects (n=13) and from patients with end-stage dilated (n=14) or ischemic (n=8) cardiomyopathy. \**P*<0.05 and *** *P*<0.001 by one-way ANOVA and Dunnett *post hoc* tests. **C**, Correlation between *MC5R* and *POMC* mRNA expression in DCM and ICM samples. Coefficients of determination (R squared; R^2^) and *P* values are presented in the graph. **D** and **E**, qPCR analysis of *MC5R* and *POMC* mRNA expression in human induced pluripotent stem cell-derived cardiomyocytes (hiPSC-CMs) that were mechanically stretched for 24 or 48 hours. n=3 individual experiments/batches of differentiation. **F** and **G**, qPCR analysis of *MC5R* and *POMC* mRNA expression in hiPSC-CMs treated with endothelin 1 (ET-1, 100 nM) for 24 hours. n=4 individual experiments/batches of differentiation. * *P*<0.05, ** *P*<0.01 versus Control by randomized block ANOVA (using individual experiments and treatment as factors). **H**, qPCR analysis of *NPPA*, *ACTA1*, *ACTA2*, *TGFB*, *CTGF* and *FN1* mRNA expression in hiPSC-CMs treated with different concentrations of PG-901 for 24 hours in the absence or presence of ET-1 (100 nM). n=4 individual experiments/batches of differentiation. * *P*<0.05 for the indicated comparisons by randomized block ANOVA (using individual experiments and treatment as factors) and Dunnett *post hoc* tests.

Since *MC5R* appeared to be expressed in hiPSC-CMs and regulated by hypertrophic stimuli, we next investigated the effects of the selective MC5-R agonist PG-901 in these cells under basal and ET-1-stimulated conditions. Of note, PG-901 reduced *TGFB1, CTGF* and *FN1* mRNA levels at 0.1 nM or 10 nM concentration in both unstimulated and stimulated hiPSC-CMs (**Figure 5H**), while *ACTA1* was downregulated at the highest tested concentration (1 μM) in ET-1-stimulated cells (**Figure 5H**). Taken together, these results demonstrate that MC5-R is functionally operative also in human cardiomyocytes.

### Cardiomyocyte-Specific MC5-R Deficiency Aggravates Pathological Cardiac Hypertrophy

To test for a regulatory role of MC5-R in cardiac hypertrophy, we engineered a tamoxifen-inducible cardiomyocyte-specific MC5-R KO mice (Mc5r-cKO) by crossing Mc5r^fl/fl^ mice with Myh6-MCM transgenic mice. Analysis of genomic DNA samples showed efficient Cre-lox recombination specifically in the heart after tamoxifen treatment that also resulted in ∼50 % reduction in cardiac MC5-R protein level (**Supplementary Figure XI**). Eight-week-old Mc5r-cKO mice and their age-matched controls (Mc5r^fl/fl^ and Myh6-MCM) were subjected to sham or TAC surgery and cardiac phenotyping was performed 4 weeks after surgery. No genotype difference was observed in sham-operated mice (**Figure 6A-D**). However, Mc5r-cKO mice manifested a subtle but significant increase in the hypertrophic response to TAC surgery compared to control Mc5r^fl/fl^ and Myh6-MCM mice (**Figure 6A-D)**. Enhanced hypertrophic response was also apparent at the cellular level, where TAC-operated Mc5r-cKO mice demonstrated increased myocyte cross-sectional area (**Figure 6A** and **6E**). Supporting these findings, echocardiographic analysis revealed enhanced thickening of the LV posterior wall in Mc5r-cKO mice after TAC surgery (**Figure 6F**). In terms of LV systolic function, TAC-operated mice did not show any deterioration of LV ejection fraction compared to sham-operated mice (**Figure 6G**). However, tracing of the LV endocardial border to measure fractional area change (FAC) revealed depressed LV systolic function in TAC-operated mice and Mc5r-cKO mice showed lower FAC compared to Myh6-MCM mice among the TAC-operated groups (**Figure 6H**). Furthermore, Mc5r-cKO mice showed reduced isovolumetric relaxation time (**Figure 6I**) and increased mitral annular e’/a’ ratio after TAC surgery (**Supplementary Figure XII**), while other parameters of diastolic function were unchanged (**Supplementary Figure XII**).

**Figure 6.**
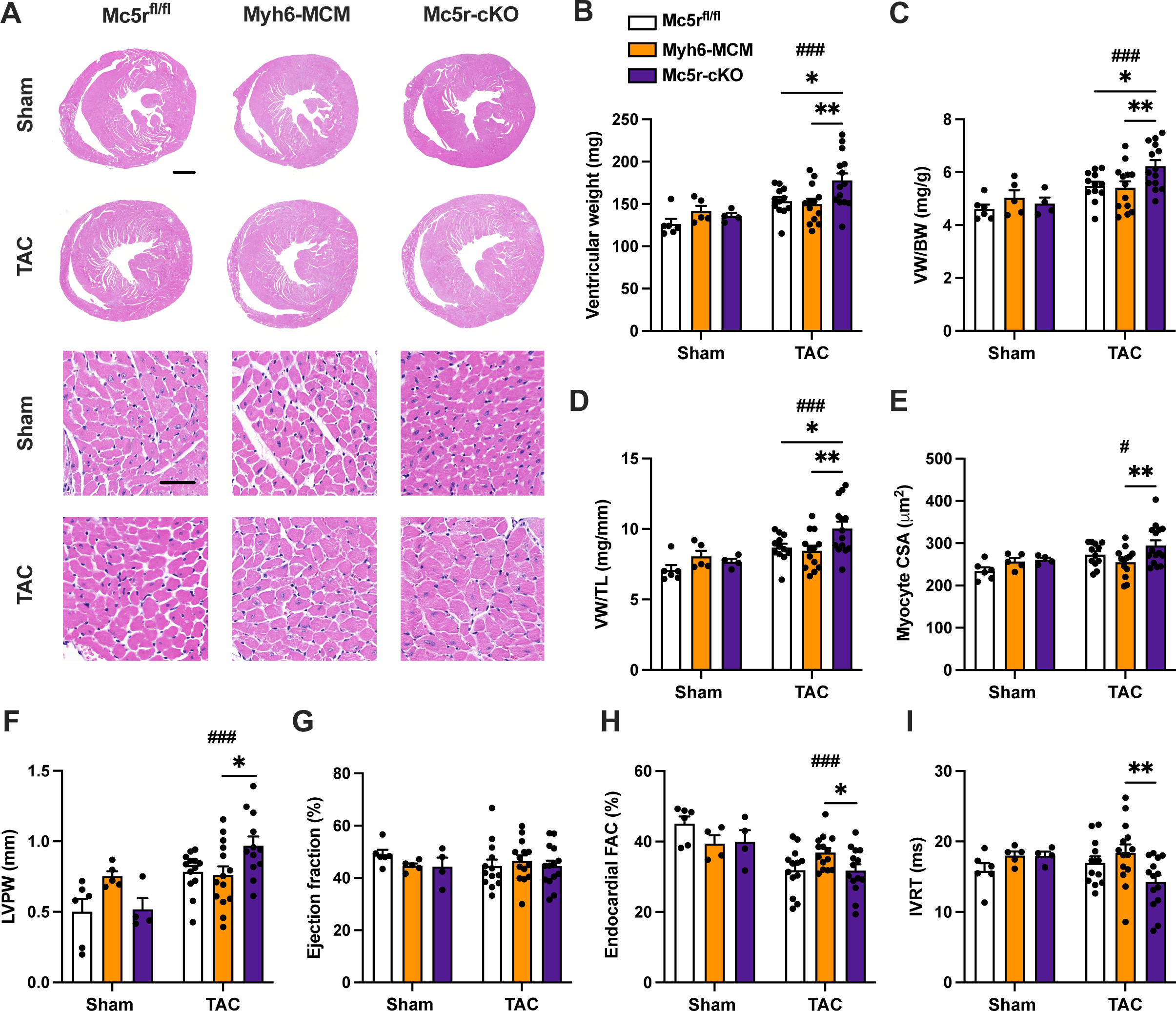
Cardiomyocyte-restricted MC5-R deficiency aggravates cardiac hypertrophy after pressure overload. **A**, Exemplary hematoxylin and eosin (H&E)-stained cross-sections of the heart showing the gross morphology of Mc5r^fl/fl^, Myh6-MCM and Mc5r-cKO mice after 4 weeks of sham or TAC operation. Scale bars, 1 mm (upper panel) and 20 μm (lower panel). **B** through **D**, Ventricular weight, ventricular weight to tibia length ratio (VW/TL) and ventricular weight to body weight ratio (VW/BW) in the indicated groups. **E**, Quantification of cross-sectional area of ventricular cardiomyocytes. **F** through **I**, Echocardiographic analysis of LV posterior wall thickness (LVPW), ejection fraction, endocardial fractional area change (FAC) and isovolumetric relaxation time (IVRT) in Mc5r^fl/fl^, Myh6-MCM and Mc5r-cKO mice after 4 weeks of sham or TAC operation. Data are mean ± SEM, each dot represents individual mouse. * *P*<0.05 and ** *P*<0.01 for the indicated comparisons by 2-way ANOVA and Dunnett *post hoc* tests. ## *P*<0.01, ### *P*<0.001 for the main effect of TAC by 2-way ANOVA.

Corroborating the findings of MC5-R evoked effects on fibrosis-associated genes *in vitro*, the extent of perivascular and interstitial fibrosis was increased in the LV of TAC-operated Mc5r-cKO mice (**Figure 7A** and **7B**). Gene expression analyses by qPCR also revealed upregulation of the hypertrophic marker genes *Nppa* and *Nppb*, fibrotic genes *Ctgf*, *Mmp2* and *Fn1* and the proinflammatory gene *Il6* in Mc5r-cKO mice after TAC surgery (**Figure 7C** through **7H**, **Supplementary Figure XII**).

**Figure 7.**
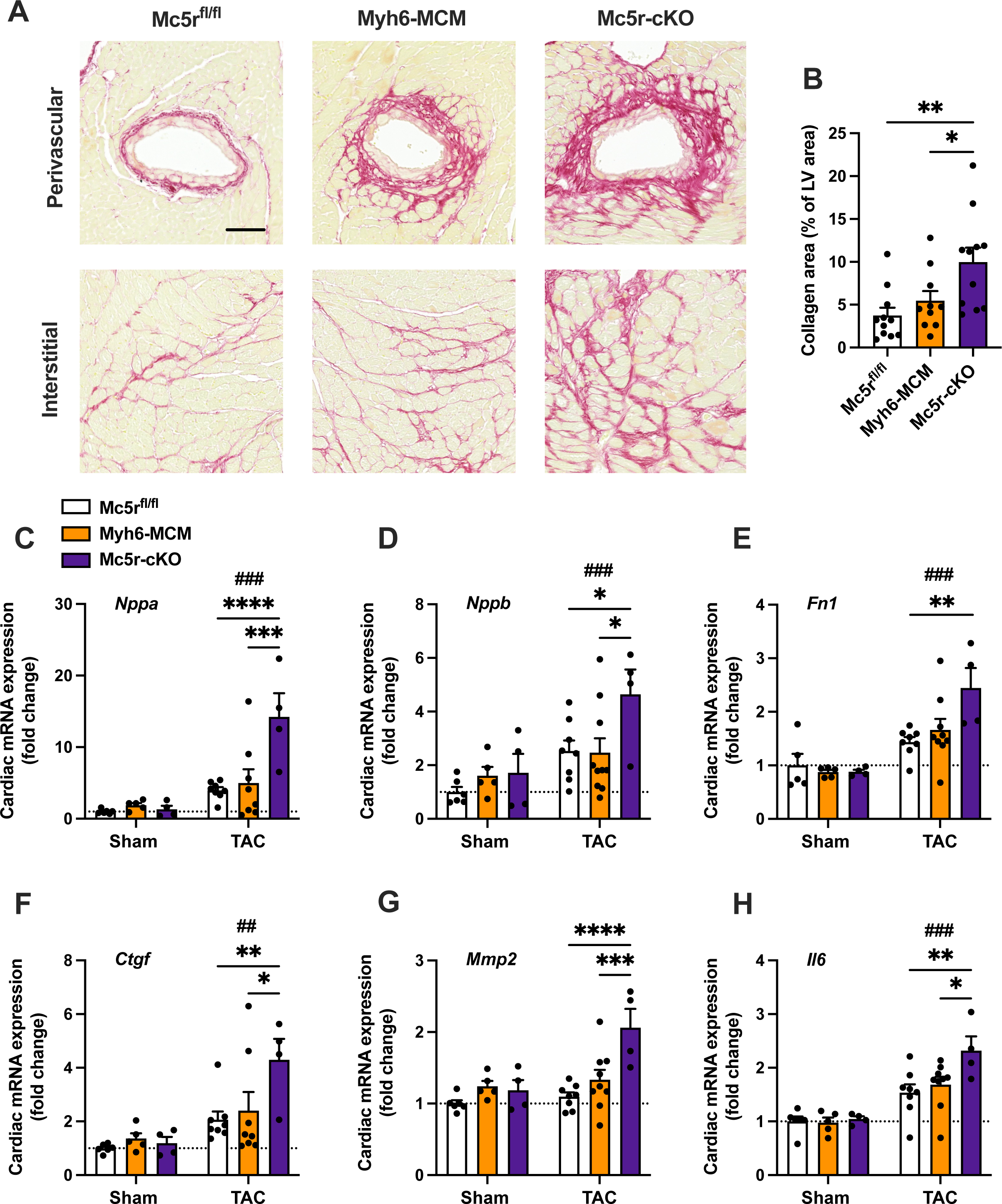
Cardiomyocyte-restricted MC5-R knockout mice show enhanced cardiac fibrosis after TAC surgery. **A**, Exemplary Picrosirius Red-stained cross-sections of the LV showing the extent of perivascular and interstitial fibrosis in Mc5r^fl/fl^, Myh6-MCM and Mc5r-cKO mice after TAC surgery. Scale bar, 100 μm. **B**, Comparison of the LV collagen area in Mc5r^fl/fl^, Myh6-MCM and Mc5r-cKO mice after TAC surgery. **C** through **H**, Quantitative real-time PCR (qPCR) analysis of *Nppa* (atrial natriuretic peptide), *Nppb* (brain natriuretic peptide), *Fn1* (fibronectin), *Ctgf* (connective tissue growth factor), *Mmp2* (matrix metalloproteinase 2) and *Il6* (interleukin 6) in the LV of Mc5r^fl/fl^, Myh6-MCM and Mc5r-cKO mice after sham or TAC surgery. Data are mean ± SEM, each dot represents individual mouse. * *P*<0.05, ** *P*<0.01, *** *P*<0.001 and **** *P*<0.0001 for the indicated comparisons by 2-way ANOVA and Dunnet *post hoc* tests. ## *P*<0.01, ### *P*<0.001 for the main effect of TAC by 2-way ANOVA.

In contrast to TAC model, four-week infusion of Ang II induced a clear hypertrophic response but Mc5r-cKO mice were not sensitized to this response compared to their control genotypes (**Supplementary Figure XIII**). Furthermore, cardiac function and structure, as assessed by echocardiography, did not reveal any genotype differences in Ang II-infused mice (**Supplementary Table IV**).

### Pharmacological Activation of MC5-R Protects against Heart Failure

Finally, to evaluate the therapeutic potential of targeting MC5-R for the management of heart failure, we employed the TAC-model in C57Bl/6N mice that are more prone to develop TAC-induced heart failure^38,39^ and treated the mice with PG-901 at two different dose levels (0.005 or 0.5 mg/kg/day). TAC-operated C57Bl/6N mice developed robust hypertrophic response in terms of ventricular weight (**Figure 8A** through **8C**), thickening of LV posterior wall (**Figure 8D**) and LV dilatation (**Supplementary Figure XIV**) but no significant treatment effect was noted for PG-901 in this regard. However, echocardiography revealed that TAC-operated mice treated with the low dose of PG-901 had a significant improvement in LV ejection fraction compared to vehicle-treated mice (**Figure 8E**). PG-901 treatment also protected against TAC-induced reduction in mitral valve deceleration time (**Figure 8F**), a measure of ventricular stiffness during diastole. Other parameters of diastolic function were unchanged in PG-901-treated mice compared to vehicle-treated TAC mice (**Supplementary Figure XIV**). The observed functional changes were associated with a reduction in the extent of LV fibrosis (**Figure 8A** and **G**). In terms of molecular features of heart failure, low dose of PG-901 downregulated cardiac expression of *Nppa*, *Mmp2* and *Ctgf* (**Figure 8H**). Taken together, these findings suggest that pharmacological targeting of MC5-R signaling could provide therapeutic benefits in the management of heart failure.

**Figure 8.**
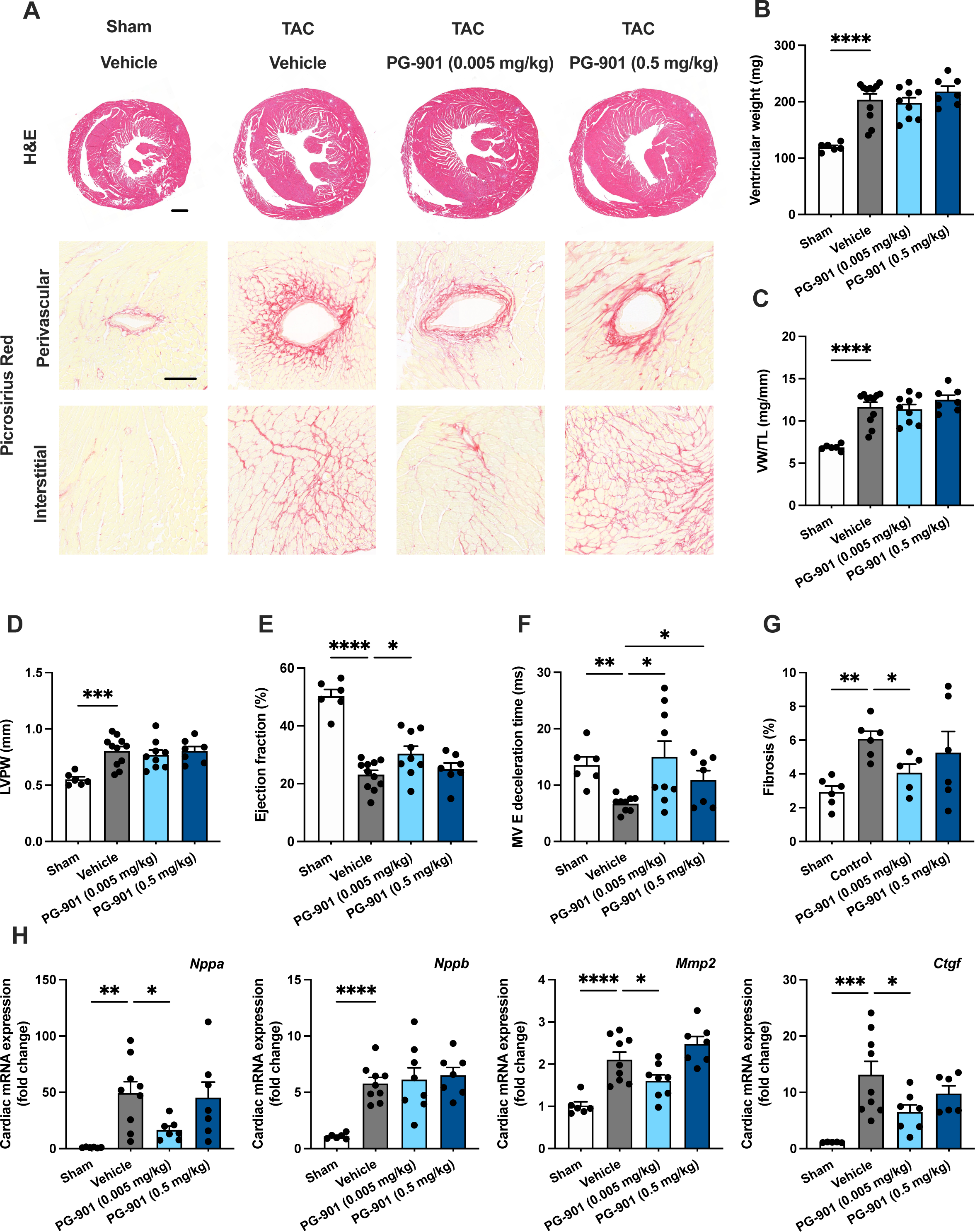
MC5-R activation improves LV systolic function and reduces cardiac fibrosis in TAC-operated mice. **A**, Exemplary hematoxylin and eosin (H&E)- and Picrosirius Red-stained cross-sections of the heart showing the gross morphology and the extent of perivascular and interstitial fibrosis in sham- and TAC-operated mice treated with either vehicle or PG-901 (0.5 or 0.005 mg/kg/day). Scale bar, 1 mm (H&E), 100 μm (Picrosirius Red). **B** and **C**, Ventricular weight and ventricular weight to tibia length ratio (VW/TL) in the indicated groups. **D** and **E**, Left ventricular posterior wall thickness (LVPW) and ejection fraction analyzed by echocardiography at the end of the experiment. **F**, Mitral valve (MV) deceleration time analyzed by pulsed-wave Doppler echocardiography at the end of the experiment. **G**, Quantification of the extent of LV fibrosis. **H**, Quantitative real-time PCR (qPCR) analysis of *Nppa* (atrial natriuretic peptide), *Nppb* (brain natriuretic peptide), *Mmp2* (matrix metalloproteinase 2) and *Ctgf* (connective tissue growth factor) in the LV of sham- and TAC-operated mice treated with either vehicle or PG-901. Data are mean ± SEM, each dot represents individual mouse. * *P*<0.05, ** *P*<0.01, *** *P*<0.001 and *P*<0.0001 for the indicated comparisons by 1-way ANOVA and Dunnett *post hoc* tests.

## Discussion

Our study uncovers a new melanocortin signaling pathway in the heart that is involved in the hypertrophic remodeling of the myocardium. First, we identified that *Pomc* and α-MSH expression in the heart is modulated by experimental pressure overload and showed that elevating α-MSH level protects against pathological cardiac hypertrophy and associated deterioration of LV systolic function. Secondly, our results establish a mechanistic link to MC5-R that is functionally active in ventricular myocytes and mediates antihypertrophic and antifibrotic response upon activation. Conversely, silencing MC5-R signaling in cardiomyocytes aggravates pressure overload-induced cardiac hypertrophy and fibrosis. Taken together, these findings provide the first evidence that a functional melanocortin circuit exists in the heart and it regulates cardiac growth response in a favorable manner.

As a first line of evidence for the involvement of α-MSH in cardiac remodeling, we observed that acute pressure overload after TAC surgery triggered a change in cardiac *Pomc* and α-MSH expression with cardiomyocytes emerging as the primary source of α-MSH. It also appeared that there is a biphasic response in cardiac *Pomc* expression during the progression of heart failure with increased number of *Pomc*^+^ - cells in compensated hypertrophy and then declining level in the failing heart. These results are in agreement with the clinical finding that plasma α-MSH level inversely correlated with NYHA functional class in heart failure patients.^15^ Considering that plasma α-MSH concentration is relatively low (∼10 pg/ml vs ∼300 pg/mg protein in the heart) and it was not changed in TAC-operated mice, it is likely that α-MSH primarily acts in an autocrine or paracrine fashion in the heart without being significantly released into the circulation. To investigate whether the declining level of α-MSH in the failing heart could be counteracted by pharmacological means to provide therapeutic benefits, TAC-operated mice were chronically treated with α-MSH analogue. Indeed, repeated α-MSH administration reduced ventricular weight and improved LV systolic function in TAC-challenged mice, demonstrating that α-MSH protects against pathological cardiac hypertrophy.

As a secreted peptide hormone, α-MSH may act on ventricular cardiac myocytes or other target cells in the heart such as fibroblasts, endothelial cells or macrophages, which are all known to express functional MC-Rs. Although we cannot exclude the involvement of other cell types as mediators of the antihypertrophic regulation of α-MSH, *in vitro* experiments with H9c2 cells and NMCMs proved that cardiomyocytes are responsive to α-MSH treatment and thus express functional MC-Rs. In the quest of the responsible MC-R subtype for the antihypertrophic regulation, we found that MC5-R activation mimics the effects of α-MSH and that the effects of α-MSH were reversed by MC5-R antagonism, supporting the notion that MC5-R is the primary receptor responsible for mediating the antihypertrophic regulation of α-MSH. Intriguingly, MC5-R is expressed in the mouse heart and the dimer form of MC5-R protein was increased in the LV of hypertrophied heart with normal ejection fraction, while in heart failure, the amount of MC5-R dimer form was significantly reduced.

Thus, the progression from compensated hypertrophy to heart failure led to parallel changes in cardiac MC5-R expression and α-MSH level, suggesting that exhaustion of α-MSH production simultaneously compromises the integrity of MC5-R. Supporting this notion, a highly significant correlation between the expression of *MC5R* and *POMC* was observed in human LV samples and hiPSC-CMs.

Of particular importance, we found that functional MC5-R is expressed in mouse and human cardiomyocytes and identified a novel role for this MC-R subtype in cardiac remodeling. Previous studies have shown that MC5-R is most abundantly expressed in the skin, adrenal gland and skeletal muscle, while low but detectable levels of MC5-R mRNA have been found in various peripheral tissues including the heart.^40-42^ However, the functional significance of MC5-R in the heart has remained unexplored. A recent study that has the closest relevance to our work reported that H9c2 cells express MC5-R and respond to treatment with α-MSH or PG-901 at subnanomolar concentrations.^43^ The authors aimed to build on the finding that α-MSH, by interacting with MC5-R, promotes glucose uptake in the skeletal muscle and expanded this concept to H9c2 cells by showing that MC5-R activation modulates the expression of glucose transporters and protects H9c2 cells against high glucose-induced apoptosis and hypertrophy.^43,44^ Our findings corroborate the existence of functional MC5-R in H9c2 cells and further prove that these cells respond to treatment with α-MSH or PG-901 in a similar way as primary cardiomyocytes. The effects of α-MSH and selective MC5-R agonism with PG-901 evoked similar responses in H9c2 cells and NMCMs but the kinetic response profiles differed slightly with PG-901 showing more sustained effects in terms of JNK inhibition and downregulation of hypertrophy- and fibrosis-associated genes. This difference could be explained by the cyclic structure of PG-901 that makes it resistant to proteolytic degradation and thus biologically more stable compared to α-MSH.^26^ PG-901 has been characterized to be a full agonist at human MC5-R with an EC_50_-value below 0.1 nM in a cAMP assay,^26^ which is consistent with the present study showing appearance of effects at subnanomolar concentrations of PG-901. The *in vitro* results were also well in line with the dose-response relationship of PG-901 in TAC-operated mice.

Screening of the intracellular signaling responses revealed that MC5-R activation reduces cAMP level, indicative of Gi coupling. It has been previously established that MC5-R can engage two parallel signaling pathways upon activation: Gs/cAMP/PKA and Gi/ERK1/2.^33^ Despite the observed reduction in cAMP level, further mechanistic experiments showed that the downstream effects are independent of the Gi pathway. We also found that the signal transduction ensuing from MC5-R activation does not rely on Gs/cAMP/PKA axis either. Nevertheless, a more profound and consistent reduction was observed on the level of phosphorylated JNK after MC5-R activation. Intracellular signaling of MC-Rs has been rarely linked to the JNK pathway and its inhibition but a recent study demonstrated that melanocortin signaling through MC5-R can inhibit JNK activity in mouse adipocytes.^45^ The exact molecular mechanism for the JNK inhibition has remained elusive, but in the case of cardiomyocytes, it appears to be driven by a cAMP-independent signaling cascade.

JNKs belong to a subclass of stress-activated protein kinases (SAPKs) and in cardiomyocytes, their activation can be promoted by GPCRs, receptor tyrosine kinases and a variety of different stress stimuli including oxidative stress and ischemia. Remarkably, *in vitro* and *in vivo* studies addressing the role of JNK signaling in cardiac hypertrophy have yielded conflicting results.^46^ Evidence obtained from *in vitro* studies strongly argue for a prohypertrophic role of JNKs, while loss-of-function approaches to silence JNK or its upstream regulators *in vivo* have resulted in promotion as well as attenuation of cardiac growth response.^47-49^ *In vitro* experiments in the current study demonstrate that MC5-R activation with PG-901 downregulates fibrosis-related genes in a JNK-dependent manner, which therapeutically translated into a clear anti-fibrotic effect in TAC-challenged mice without suppression of the hypertrophic response. These findings might relate to the inconclusive studies on the functional role of JNKs in the cardiac hypertrophic response, and suggest that JNKs have a more significant role as regulators of cardiac fibrosis. However, further studies are warranted to determine whether MC5-R-induced JNK reduction has a direct effect on cardiomyocyte growth *in vivo*.

Corroborating the regulatory role of MC5-R in cardiac remodeling, our loss-of-function study revealed that silencing MC5-R specifically in cardiomyocytes renders mice more susceptible to TAC-induced cardiac hypertrophy and fibrosis. The cardiac phenotype of Mc5r-cKO mice matched closely the phenotype of cardiomyocytes with siRNA-induced knockdown of *Mc5r*. *In vitro* and *in vivo* silencing of MC5-R were both associated with upregulation of *Nppa*, *Nppb*, *Ctgf* and *Fn1*. Conversely, MC5-R activation *in vivo* improved LV systolic function in TAC-operated mice and it was accompanied by reduced cardiac fibrosis and downregulation of *Nppa* and *Ctgf*. CTGF, for instance, is strongly produced by injured cardiomyocytes and it regulates many fibrosis-related processes such as extracellular matrix deposition.^50,51^ CTGF also stimulates hypertrophic growth of cultured cardiomyocytes.^52,53^ Given the involvement of MC5-R in cardiac fibrosis, it will be intriguing to further explore whether functional MC5-R exists also in cardiac fibroblasts and could it synergize with MC5-R in cardiomyocytes to regulate remodeling of the myocardium. On the other hand, fibroblasts are considered as important effector cells in the hypertrophic response to Ang II and Mc5r-cKO mice were not sensitized to Ang II-induced cardiac hypertrophy.^54,55^ Furthermore, Ang II infusion did not change MC5-R expression in the heart, while TAC surgery clearly affected cardiac MC5-R protein levels. These findings suggest that MC5-R signaling does not modulate hypertrophic remodeling that is primarily driven by cardiac fibroblasts or other non-myocytes.

As a limitation for drawing conclusion on the significance of MC5-R in cardiac remodeling, tamoxifen-induced Mc5r-cKO mice developed only a mild increase in ventricular weight after 4 weeks of TAC surgery compared to control mice. In order to avoid Cre-mediated cardiotoxicity, tamoxifen dosing needs to be lowered and ideally fractionated by administering tamoxifen on consecutive days.^17,56^ This however brings along the problem of insufficient recombination and gene knockdown, which might be the case also in the current study, since Mc5r-cKO mice showed only a partial reduction (∼50 %) of MC5-R expression in the heart. This, in turn, might explain the subtle phenotype of Mc5r-cKO mice and underestimate the role of MC5-R in cardiac remodeling. Another contributing factor might be that TAC-induced pressure overload was found to reduce the dimer form of MC5-R, thus limiting the incremental effect of genetically-induced MC5-R deficiency on the hypertrophic response.

In conclusion, the present study uncovers a novel role for α-MSH and MC5-R in pathological cardiac remodeling. α-MSH is expressed in the heart and protects against pathological cardiac hypertrophy by activating MC5-R in cardiac myocytes, which may be a potential therapeutic target for the management of heart failure. Considering that analogues of naturally occurring α-MSH have been recently approved for clinical use (Scenesse®, Vyleesi®, and Imcivree®),^57^ it is important to further evaluate the cardiovascular safety of melanocortin drugs. Even if future research does not support the translation of MC5-R targeted drugs for treating human heart failure, the present findings predict a favorable cardiac safety profile for drugs that have agonistic activity at MC5-R.

## Acknowledgements

We thank Sanna Bastman, Hanna Haukkala, Satu Mäkelä, Johanna Jukkala and Annika Korvenpää for their excellent technical support. The histological methods were performed by the Histology core facility of the Institute of Biomedicine, University of Turku, Finland.

## Novelty and Significance

### What Is Known?

- α-Melanocyte-stimulating hormone (α-MSH), primarily expressed in the pituitary gland and skin, regulates several physiological functions by interacting with melanocortin receptors (MC1-R – MC5-R)
- α-MSH has been previously shown to be also expressed in the rat heart and its plasma levels are changed in patients with cardiomyopathies.
- However, the role of α-MSH and its possible target receptors in the heart has remained elusive.

### What New Information Does This Article Contribute?

- The production of α-MSH in the heart declines in the failing mouse heart and pharmacological administration of α-MSH protects against cardiac hypertrophy induced by pressure overload.
- The protective effects of α-MSH are mediated by the melanocortin 5 receptor (MC5-R), which is expressed in mouse and human cardiac myocytes.
- Silencing of MC5-R in cardiomyocytes renders mice susceptible for enhanced cardiac hypertrophy and fibrosis after pressure overload.

α-MSH is a pleiotropic peptide hormone that is proteolytically cleaved from the precursor molecule pro-opiomelanocortin (POMC). It is predominantly produced by the pituitary gland, hypothalamic neurons and melanocytes in the skin. Earlier evidence demonstrates its presence also in the heart, but the functional significance of α-MSH in the heart is completely unknown. In this study, we found that cardiac *Pomc* and α-MSH expression declines when the heart undergoes maladaptive remodeling after pressure overload. Pharmacological treatment of pressure overloaded mice with α-MSH attenuated cardiac hypertrophy and systolic dysfunction. Mechanistically, α-MSH downregulated hypertrophy- and fibrosis-associated genes and reduced cellular growth of cardiomyocytes by activating melanocortin 5 receptor (MC5-R). In contrast, MC5-R deficiency in cardiomyocytes promoted cardiac hypertrophy and fibrosis in mice. The present findings suggest that α-MSH or more specific activation of MC5-R may translate into therapeutic benefits in the management of pathological cardiac remodeling and heart failure.

